# Novel role for ESCRT-III component CHMP4C in the integrity of the endocytic network utilized for herpes simplex virus envelopment

**DOI:** 10.1101/2020.08.19.258558

**Authors:** Tiffany Russell, Jerzy Samolej, Michael Hollinshead, Geoffrey L. Smith, Joanne Kite, Gillian Elliott

## Abstract

Enveloped viruses exploit cellular trafficking pathways for their morphogenesis, providing potential scope for the development of new antiviral therapies. We have previously shown that herpes simplex virus 1 (HSV1) utilises recycling endocytic membranes as the source of its envelope, in a process involving four Rab GTPases. To identify novel factors involved in HSV1 envelopment, we have screened an siRNA library targeting over eighty human trafficking proteins including coat proteins, adaptor proteins, fusion factors, fission factors and Rab effectors. Depletion of eleven factors reduced virus yield by 20- to 100-fold, including three early secretory pathway proteins; four late secretory pathway proteins; and four endocytic pathway proteins, three of which are membrane fission factors. Five of the eleven targets were chosen for further analysis in virus infection where it was found that the absence of only one, the fission factor CHMP4C, known for its role in the cytokinesis checkpoint, specifically reduced virus production at the final stage of morphogenesis. Ultrastructural and confocal microscopy of CHMP4C-depleted, HSV1-infected cells, showed an accumulation of endocytic membranes; extensive tubulation of recycling, transferrin receptor-positive endosomes indicative of aberrant fission; and a failure in virus envelopment. No effect on the late endocytic pathway was detected, while exogenous CHMP4C was shown to localise to recycling endosomes. Taken together, these data reveal a novel role for the CHMP4C fission factor in the integrity of the recycling endosomal network, which has been unveiled through the dependence of HSV1 on these membranes for the acquisition of their envelopes.

**Importance:** Cellular transport pathways play a fundamental role in secretion and membrane biogenesis. Enveloped viruses exploit these pathways to direct their membrane proteins to sites of envelopment, and as such, are powerful tools for unravelling subtle activities of trafficking factors, potentially pinpointing therapeutic targets. Using the sensitive biological readout of virus production, over eighty trafficking factors involved in diverse and poorly defined cellular processes have been screened for involvement in the complex process of HSV1 envelopment. Out of eleven potential targets, CHMP4C – a key component in the cell-cycle abscission checkpoint – stood out as being required for the physical process of virus wrapping in endocytic tubules, where it was shown to localise. In the absence of CHMP4C, recycling endocytic membranes failed to undergo scission, causing transient tubulation and accumulation of membranes and unwrapped virus. These data reveal a new role for this important cellular factor in the biogenesis of recycling endocytic membranes.

## Introduction

During the process of virus envelopment, intracellular trafficking pathways are subverted to supply lipid membranes for virus wrapping and release. These membranes contain virus-encoded glycoproteins that play a role in attachment and fusion during virus entry. While all virus glycoproteins start their life in the endoplasmic reticulum (ER), the cellular sites to which they are subsequently transported and where envelopment occurs vary between virus families [1]. Herpes simplex virus type 1 (HSV1) is a large enveloped virus that has an intricate morphogenesis pathway termed the envelopment-deenvelopment-reenvelopment pathway [2, 3] in which capsids form in the nucleus and bud through the inner nuclear membrane as primary virions [4–6], using virus-encoded machinery termed the nuclear egress complex [7]. The primary envelope is lost by fusion with the outer nuclear membrane, releasing naked capsids into the cytosol [4, 5]. As for cellular membrane proteins, HSV1 envelope glycoproteins are co-translationally inserted into the ER and transported through the secretory pathway to the Golgi apparatus and plasma membrane [8], with free capsids acquiring their final envelope from a wrapping site within the cytoplasm that contains these glycoproteins. Previously, we defined a model in which HSV1 acquires its envelope from glycoprotein-containing endocytic membranes that have been recently retrieved from the plasma membrane [9] rather than the TGN as often cited [10–12], and in agreement with earlier studies from others [13]. In this model, virus egress would then occur through the natural recycling of these membranes to the cell surface. This model has been supported by more recent studies showing that glycoproteins must be transported to the plasma membrane and endocytosed prior to envelopment taking place [14, 15].

Using targeted siRNA screening, four members of the Rab family of GTPases were identified as important for HSV1 envelopment, including Rab1 in the early secretory pathway and Rab5 and Rab11 in the endocytic pathway [9, 15]. Moreover, in one of the first demonstrations of a biological role, Rab6A was shown to be critical for the transport of virus glycoproteins from the Golgi apparatus to the plasma membrane, thereby providing a source of envelope proteins for subsequent endocytic retrieval and wrapping [15]. Rab GTPases function as central regulators of the four major steps of intracellular membrane traffic - vesicle budding, delivery, tethering, and fusion - to coordinate transport and delivery pathways [16, 17]. However, they do not work alone, as these steps involve multiple cellular factors that organise, recruit, provide direction and target specific cargoes throughout the cell. The aim of this current study was to pinpoint steps along the cellular transport pathways that are exploited by HSV1 to target its envelope proteins to the correct assembly sites, where intervention would be debilitating to the virus and potentially tractable to antiviral intervention. As such, we have screened an siRNA library targeted at a rationally selected set of eighty-two factors including coat proteins involved in membrane curvature and budding [18]; adaptor proteins that select cargo for vesicles [19]; membrane fusion and fission factors [20]; and Rab effector proteins which provide specificity in vesicle targeting [21]. Eleven factors were identified whose depletion altered virus production greater than 20-fold, and which are spread across the early and late secretory pathway and the endocytic pathway. While several of these block virus infection at early stages of virus life cycle prior to morphogenesis, one of these factors – the ESCRTIII component CHMP4C – is shown to localise to and be required for the integrity of the recycling endocytic network needed for envelopment of HSV1, suggestive of a role for this protein in the scission of these endocytic membranes. Hence, this study has unveiled a novel function of CHMP4C in the cell, reinforcing the power of using virus morphogenesis to identify novel activities in cellular trafficking.

## Results

### Targeted siRNA screening identifies host trafficking factors involved in HSV1 replication

Previously, siRNA library screening was used to identify three Rab GTPases involved in HSV1 envelopment [9, 15]. To identify additional human cellular transport factors important for this process, we carried out a targeted siRNA screen directed against a range of host membrane trafficking factors which included coat proteins, adaptor proteins, fusion factors, fission factors and Rab effectors (Table S1). Because the efficiency of library knockdown was anticipated to be between 70 and 85%, two individual sets of siRNAs against the 82 targets were used in independent screening experiments to reduce the chance of missing positive targets. The siRNAs were reverse transfected into HeLa cells for 48 h, before infection with HSV1. After 24 h infection, the released extracellular virus was harvested from the medium, and quantified by plaque assay to compare the resulting yield of virus in siRNA-depleted cells to cells transfected with the negative siRNA control (blue in Fig. 1A & 1B). Depletion of the nectin1 HSV1 entry receptor (purple in Fig. 1A & 1B) [22], or the Rab GTPase RAB6A (red in Fig. 1A & 1B), which was shown to be critical for optimal HSV1 morphogenesis [15], served as positive controls. The average of three independent screens (Tables S2 & S3) revealed that the production of HSV1 was unaffected by the depletion of the majority of these host cell factors, but that six siRNAs in set 1 and eight siRNAs in set 2 reduced virus yield by 20- to 100-fold (green and orange in Fig. 1A & 1B), with three of them shared across the two sets. The identity of these eleven potential hits revealed three in the early secretory pathway (COPG1, COPG2, GOLGA2), four in the late secretory pathway (AP1B1, AP4E2, VAMP4, syntaxin 10) and four in the endocytic pathway (dynamin 2, clathrin light chain A, CHMP4C and CHMP2A) (Table 1). Interestingly, three of the four endocytic proteins were fission factors involved in membrane abscission.

**Figure 1.**
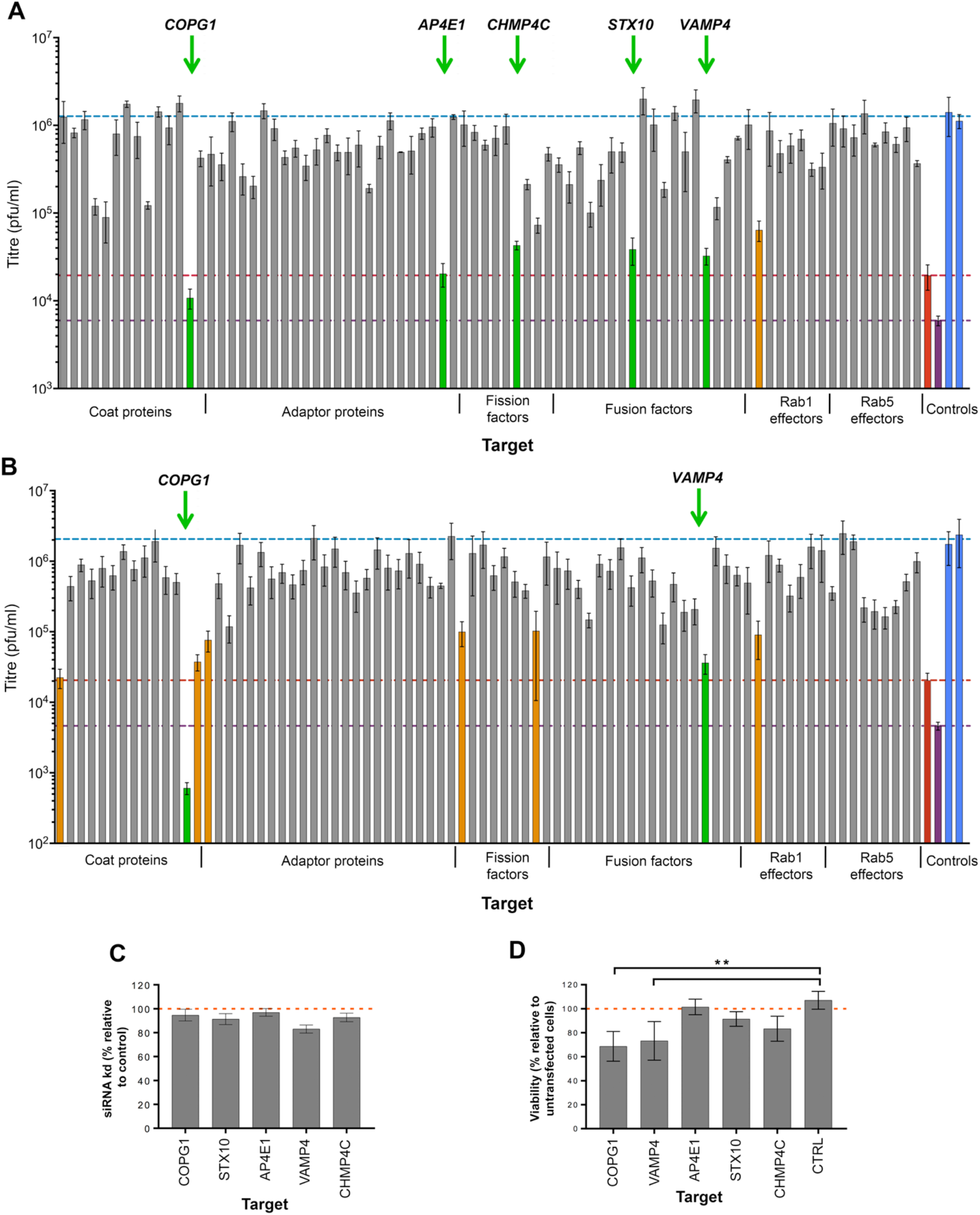
HSV1 replication in HeLa cells transfected with siRNA libraries against human trafficking proteins. (**A & B**) 20 nM of each siRNA from set 1 (**A**) and set 2 (**B**) was reverse transfected into HeLa cells in 96 well plates, incubated for 48 h and then infected with HSV1 SC16 at MOI 5 for 1 h. A gentle citrate buffer acid wash was performed to inactivate virus that had not penetrated the cells, before another 23 h of infection. Extracellular virus was harvested and titrated on Vero cells. The mean±SEM is shown, where *n* = 3. Blue – negative control; purple – nectin 1; red – Rab6A; green – factors followed up in this study; orange – factors whose depletion caused > 20-fold reduction in titre, but not followed up here. (**C**) siRNA transfection (set 1) was performed as standard. After 48 h, cells were harvested for RNA isolation, and the % siRNA knockdown (kd) was measured by RT-qPCR for the indicated targets (mean±SEM, *n* = 3). (**D**) siRNA transfection was performed as standard. After 48 h, cell viability was measured by Cell Titre Glo assay, with viability expressed as a percentage relative to untransfected cells (mean±SEM, *n* = 3).

**Table 1:** Trafficking proteins implicated in HSV1 infection using siRNA libraries.

| Target | Relative yield | Fold Drop | siRNA Set | Class | Function |
| --- | --- | --- | --- | --- | --- |
| CLTA | 0.01 | 100 | 2 | coat | Endocytosis<br>Cytokinesis |
| COPG1 | > 0.01 | 100 | 1,2 | coat | Golgi-to-ER transport |
| COPG2 | 0.02 | 50 | 2 | coat | Golgi-to-ER transport |
| AP1B1 | 0.04 | 25 | 2 | adaptor | Transport from TGN |
| AP4E1 | 0.03 | 33 | 1 | adaptor | Transport from TGN |
| DNM2 | 0.05 | 20 | 2 | fission | Endocytosis<br>Cytokinesis |
| CHMP4C | 0.03 | 33 | 1 | fission | Endosomal sorting<br>Cytokinesis |
| CHMP2A | 0.05 | 20 | 1,2 | fission | Endosomal sorting<br>Cytokinesis |
| STX10 | 0.03 | 33 | 1 | fusion | Endosome-to-TGN transport |
| VAMP4 | 0.02 | 50 | 1,2 | fusion | Endosome-to-TGN transport |
| GOLGA2 | 0.04 | 25 | 1,2 | Rab 1<br>effector | Transport through Golgi |

Five of these sixteen targets (in green, Fig. 1A) were chosen for further analysis based on their greater effect on virus yield and their involvement in diverse stages of protein trafficking and membrane remodelling: COPG1, a subunit of the COP1 coatamer complex which coats vesicles moving retrograde between Golgi stacks and from Golgi to ER [23]; AP4E1, a component of the AP4 adaptor complex found on some, non-clathrin coated vesicles leaving the trans-Golgi network (TGN) [24, 25]; syntaxin10 (STX10) and VAMP4, SNARE proteins involved in vesicle fusion with target membranes [26, 27]; and CHMP4C, a core component of the endosomal sorting required for transport complex (ESCRT-III) machinery, whose only definitive function to date is at the final stages of cytokinesis [28]. RT-qPCR was used to confirm that for all five targets of interest, the knockdown from the first set of siRNAs was greater than 80% at the transcript level (Fig. 1C). Given their potential importance for the cell, and to ensure that the requirement for each of these targets during virus infection was not a consequence of a pleiotropic and indirect effect of their depletion, a cell viability assay was also performed on depleted cells. This revealed that only depletion of COPG1 or VAMP4 resulted in a significant reduction in cell viability (Fig. 1D), and of these two, only COPG1-depleted cells showed obvious signs of morphological changes (not shown), in line with results from a previous study [29].

To determine the effect of the depletion of each of these factors on the global appearance of the secretory and endocytic pathways, siRNA transfected HeLa cells were co-stained for the Golgi marker giantin and the transferrin receptor (CD71) marker for recycling endocytic membranes. Compared to control cells, the integrity of the Golgi was compromised in many of the COPG1-depleted cells as might be anticipated from the role of COPG1 in retrograde vesicle transport through the Golgi (Fig. 2A, giantin staining). Likewise, the depletion of VAMP4 resulted in Golgi fragmentation, while the depletion of AP4E1, STX10 or CHMP4C had little effect on the steady-state appearance of the Golgi. Intracellular staining for transferrin receptor showed that as for the Golgi, the recycling endosomal network was compromised in COPG1-depleted cells (Fig. 2A, CD71 staining), and depletion of VAMP4 resulted in altered transferrin receptor localization throughout the cytoplasm. In cells depleted of the other three target factors, there was a noticeable increase in concentration of CD71 next to the nucleus, implying accumulation at the endocytic recycling compartment which localises around the microtubule organising centre (MTOC) [30].

**Figure 2.**
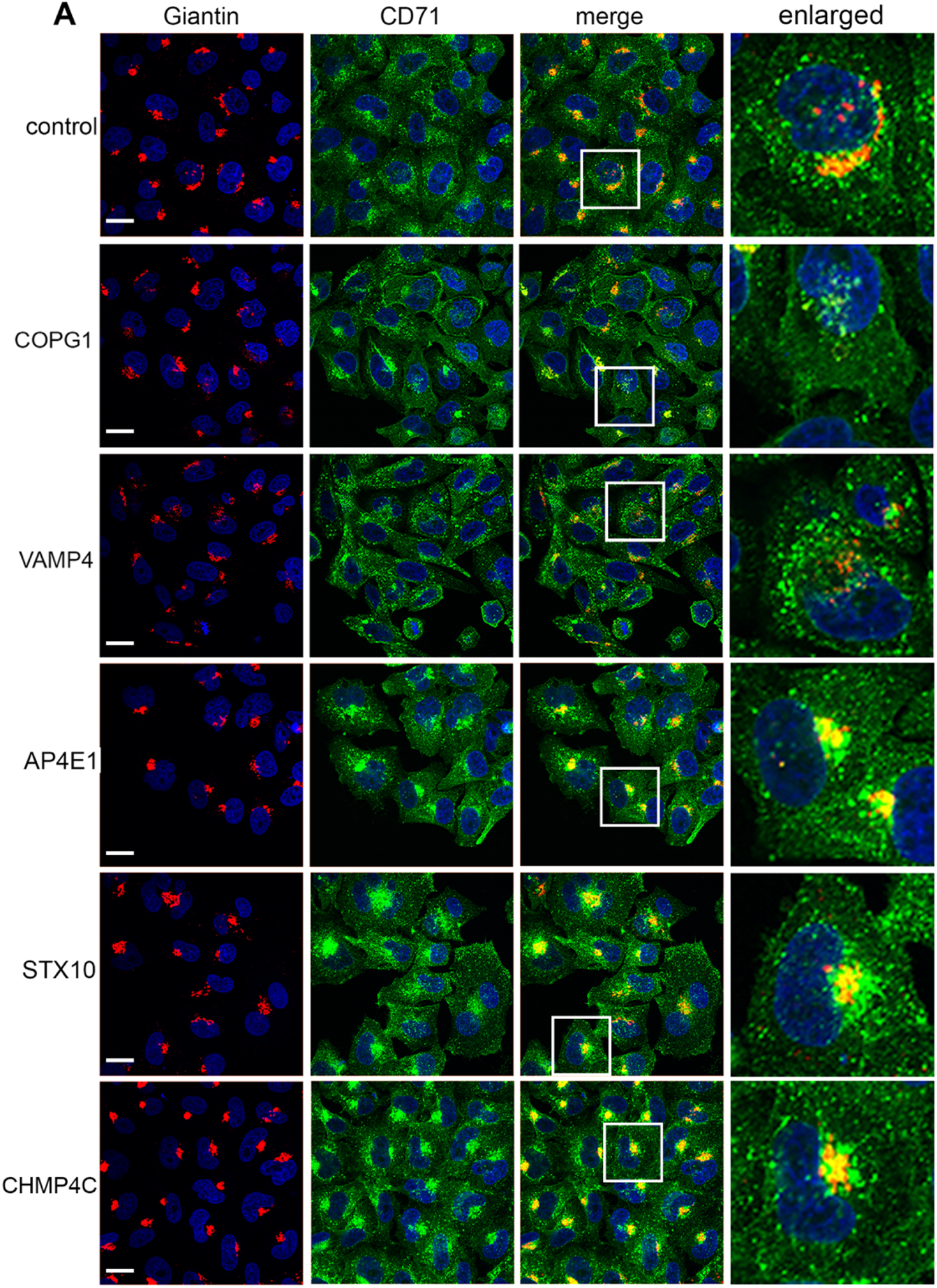
Integrity of secretory and endocytic pathways following siRNA depletion of target trafficking factors. HeLa cells transfected with negative siRNA (control) or siRNAs for COPG1, VAMP4, AP4E1, STX10 and CHMP4C were fixed after 48 h, permeabilised and stained with antibodies for the Golgi (giantin in red) and transferrin receptor (CD71 in green). Nuclei were stained with DAPI (Blue). Scale bar = 20 μm.

### Delineating the stages of virus infection affected by trafficking factor depletion

To establish where each of these identified factors was acting in the virus life cycle, we first determined if HSV1 entered cells with the equivalent efficiency after depletion of each of the targets. To measure virus entry, reverse transfection of siRNAs was performed, and after 48 h cells were infected synchronously with HSV1 expressing *β-galactosidase* under the control of the immediate early (IE) ICP0 promoter (Fig. 3A) [31]. After 4 h, *β-galactosidase* activity was measured as a surrogate for virus entry [32], although strictly, this readout could be affected by any of the steps up to IE gene expression (virus binding, membrane fusion, delivery of capsids to nucleus, and transcription of IE genes). Strikingly, depletion of COPG1 significantly impaired expression from the ICP0 promoter, to a similar level as depletion of the entry receptor nectin1, suggesting that COPG1-depleted cells may be defective for virus entry or one of the aforementioned steps. Of the other four, only CHMP4C depleted cells were able to support IE gene expression to the same level as control cells, with STX10 and AP4E1 depleted cells expressing around 2-fold lower *β-galactosidase* than control cells, and VAMP4-depleted cells reduced around 3-fold (Fig. 3A). Considering the potential for involvement of these factors in the delivery of the nectin1 receptor to the cell surface, and therefore an indirect involvement in virus entry, the effect of their depletion on localisation of nectin1 at the plasma membrane was tested. The available nectin1 antibodies were not sensitive enough to detect endogenous protein in HeLa cells, and so a plasmid expressing nectin1 tagged at its C-terminus with the V5 epitope was first constructed and tested for its expression by transient transfection of HeLa cells (Fig. S1). Western blotting indicated that nectin1-V5 was expressed as a range of high molecular weight forms (Fig. S1A), which resolved to a lower molecular weight doublet following PNGaseF treatment (Fig. S1B), indicating that nectin1 was multiply glycosylated as expected from its eight predicted N-glycosylation sites [33]. Moreover, immunofluorescence of non-permeabilised, nectin1-V5-expressing cells with the anti-nectin1 antibody (Fig. S1C) confirmed efficient cell-surface localisation of nectin1-V5. Staining of non-permeabilised, nectin-V5 expressing cells that had been transfected with siRNAs to the five targets, indicated efficient cell surface localisation of nectin1 in all except COPG1-depleted cells (Fig. 3B). However, Western blotting of transfected cell extracts revealed that this low level of cell surface nectin1 in the absence of COPG1 correlated with overall reduced nectin1 expression, further indicating the limited integrity of cells depleted for COPG1 (Fig. 3C, COPG1). Interestingly, nectin1 expressed in the absence of VAMP4 was poorly glycosylated (Fig. 3C, VAMP4), suggesting that although the depletion of VAMP4 did not affect its transport to the plasma membrane, the structure of nectin1 encountered by the virus upon binding the cell would be altered, providing a possible explanation for the apparent reduction in virus entry in VAMP4-depleted cells.

**Figure 3.**
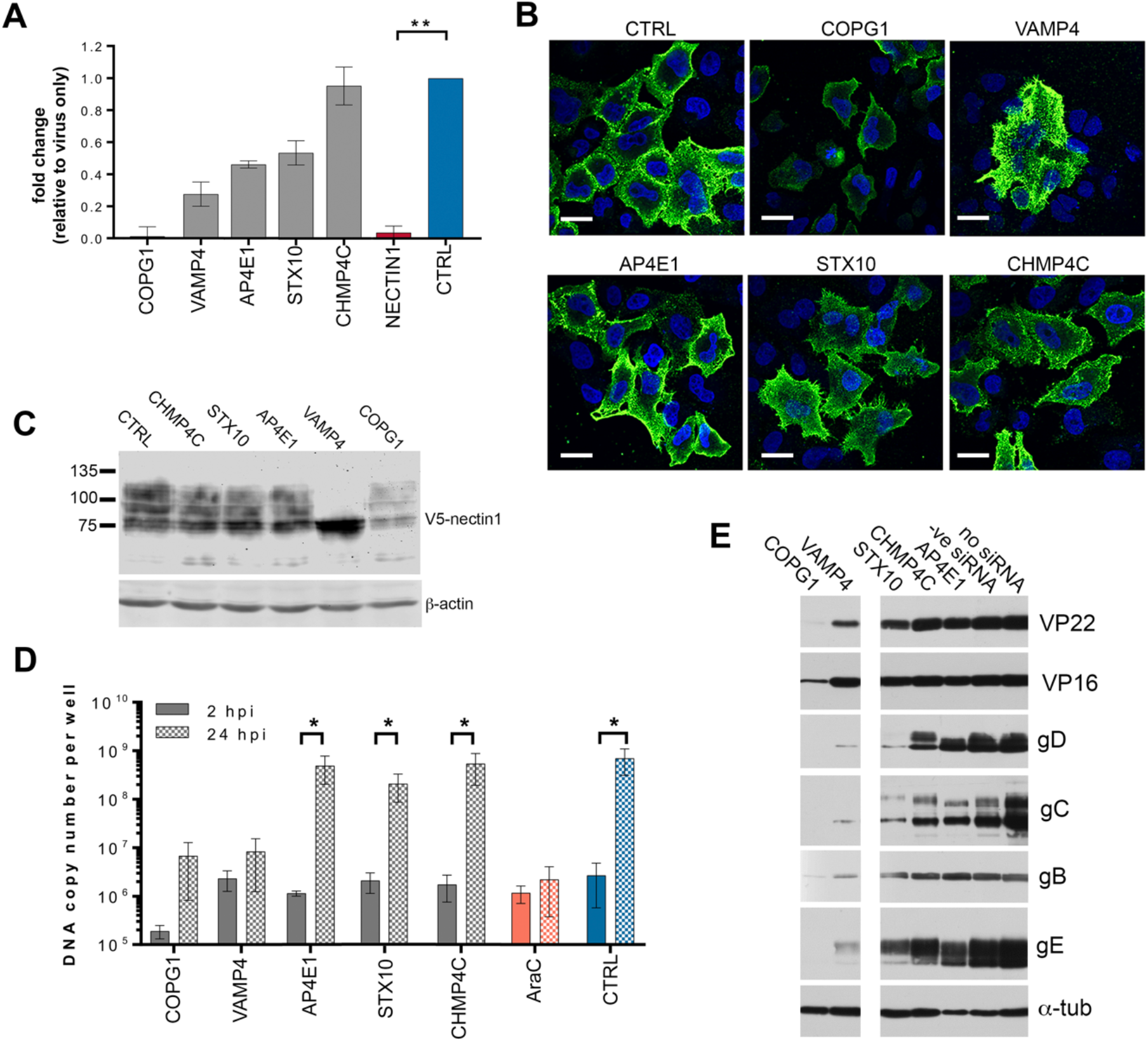
Depletion of target trafficking factors attenuates HSV1 replication at different stages of the virus life cycle. (**A**) HeLa cells transfected with siRNAs against five targets were infected 48 h later at MOI 5 with HSV1 IIE10*lacZ* for 4 h, and then β-gal activity was measured by ONPG assay (mean±SEM, *n* = 3). (**B**) HeLa cells were transfected with siRNAs against five targets together with plasmid expressing V5-tagged nectin1. After 48 h, cells were fixed and were stained without permeabilisation with CK6 antibody to detect cell surface nectin1 (green) and nuclei stained with DAPI (blue). Scale bar = 20 μm. (**C**) As for (**B**), but cells were harvested and analysed by SDS-PAGE and Western blotting for nectin 1-V5 and β-actin. (**D**) siRNA kd was performed, then cells were infected with HSV1 Sc16 at an MOI of 2. DNA was isolated at 2 h or 24 h and absolute quantitative qPCR performed for gene *UL48* to determine virus DNA copy number (mean±SEM, *n* = 3). (**E**) siRNA kd was performed, then cells were infected with HSV1 Sc16 at an MOI of 5 for 16 h before harvesting and analysing by SDS-PAGE and Western blotting for indicated proteins.

Next, the efficiency of viral DNA replication in cells transfected with siRNAs for each of our targets was compared to negative control siRNA transfection, after infection with HSV1 (Fig. 3D). Samples were harvested at 2 or 24 h after infection, and viral DNA copy number was determined by absolute qPCR for virus gene *UL48* to reflect input viral DNA or viral DNA replication, respectively. Cells were also treated with the inhibitor of HSV1 viral DNA synthesis, AraC, to serve as a positive control for inhibition of DNA replication. Depletion of AP4E1, STX10 or CHMP4C had no significant effect on DNA replication compared to the negative control, with a 1000-fold increase in DNA over the course of infection, indicating that despite some reduction in very early events of virus infection, depletion of these factors had little effect on genome replication. By contrast, depletion of VAMP4 almost abolished DNA replication, while in COPG1-depleted cells even input virus was reduced 10-fold, indicating that the greatly reduced ability of the virus to initiate immediate-early gene expression was due to reduced entry of virus genomes into the cell. Nonetheless, a modest 40-fold increase in viral DNA between 2 and 24 hpi suggests limited viral DNA replication in those cells that were able to support infection. These DNA replication results were reflected in Western blotting for late proteins in siRNA-depleted, infected cell lysates (Fig. 3E). In line with our entry and DNA replication studies, few of the virus proteins tested were detectable in COPG1-depleted cells, confirming that this infection had been blocked at very early stages of infection. VAMP4-depleted cells expressed detectable but relatively low levels of glycoproteins gB, gC, gD and gE, likely reflecting the lack of DNA replication seen in these cells. By contrast, all proteins tested were expressed to the approximate levels of control cells in CHMP4C- and AP4E1-depleted cells, confirming that the virus replication cycle was blocked late in infection in the absence of these factors. In the case of STX10-depleted cells, there was also evidence for reduced glycoprotein expression despite a normal level of DNA replication and expression of the tegument proteins VP22 and VP16 (Fig. 3E), indicating a complex phenotype of virus infection in these cells.

### Depletion of CHMP4C or STX10 alters the trafficking of recycling endosomes

As the goal of this study was to identify factors involved in HSV1 morphogenesis, only AP4E1, STX10 and CHMP4C were taken forward from this stage. It was shown previously that HSV1 glycoproteins are transported to the cell surface prior to retrieval into wrapping membranes for virus envelopment [9, 15]. Hence, to determine if glycoproteins can be delivered efficiently to the plasma membrane in AP4E1, STX10 and CHMP4C depleted cells, the cell surface localisation of glycoproteins gD and gE was examined by immunofluorescence of non-permeabilised cells at 12 hpi after infection in depleted and control siRNA transfected cells. The cell surface level of both glycoproteins was demonstrably lower in both AP4E1 and STX10-depleted cells compared to the control (Fig. 4A), although it should be noted that for STX10, this is likely due to the overall reduction in glycoprotein synthesis as shown above (Fig. 3E). By contrast, the levels in CHMP4C-depleted cells were comparable to control cells, suggesting that depletion of CHMP4C inhibits virus production at a stage downstream of glycoprotein trafficking to the cell surface.

**Figure 4.**
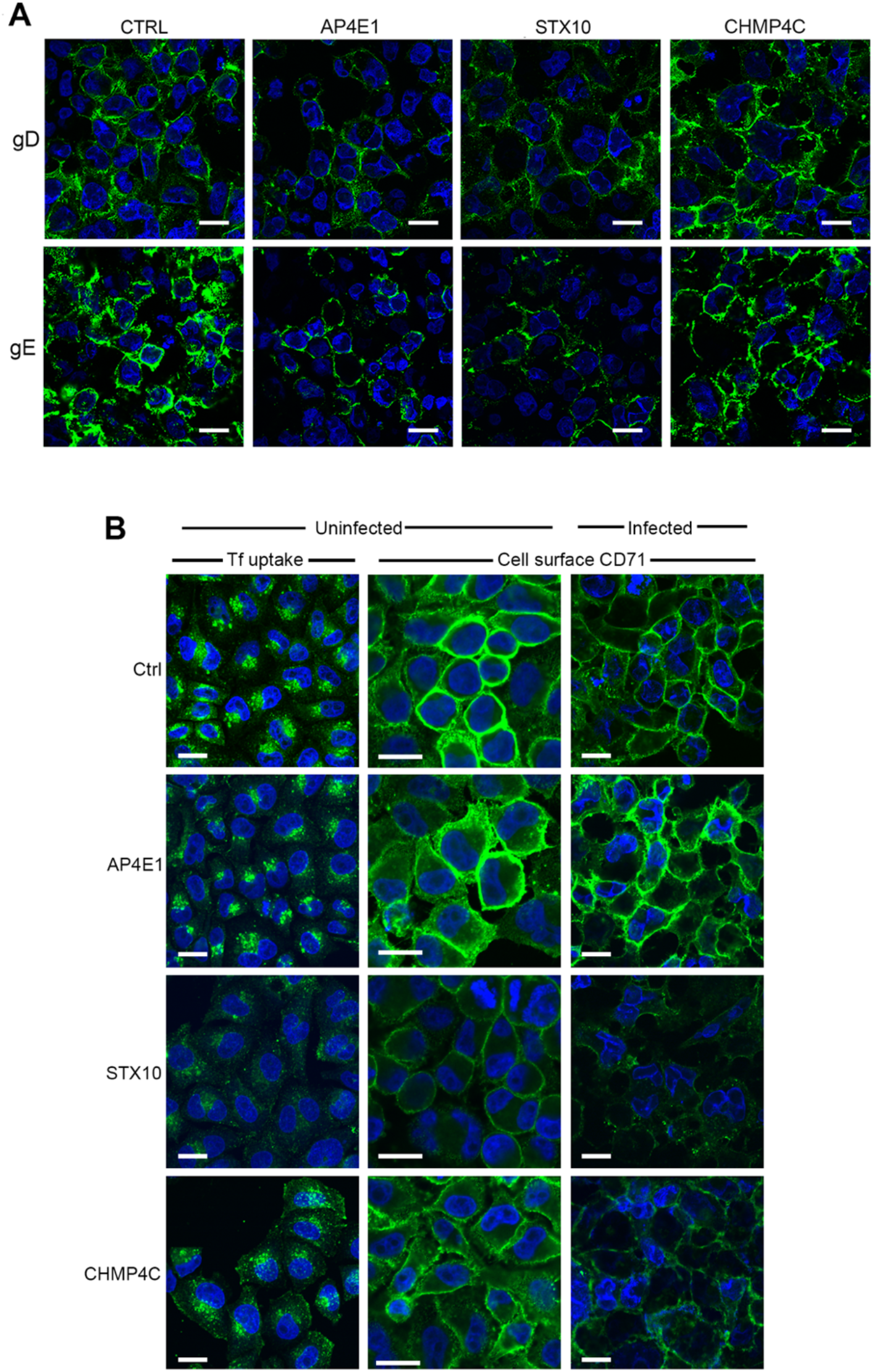
Trafficking to the cell surface in siRNA depleted cells. (**A**) siRNA knockdown was performed as indicated, then cells were infected with HSV1 Sc16 at an MOI of 5 for 12 h before fixing and cell surface staining carried out for glycoproteins D and E (green) and staining nuclei with DAPI (blue). Scale bar = 20 μm. (**B**) siRNA knockdown was performed as indicated. After 48 h cells were incubated with 488 transferrin for 30 min and fixed (Tf uptake), or were fixed in the absence of transferrin and cell-surface staining carried out for the transferrin receptor CD71 (uninfected and infected). Scale bar = 20 μm.

To determine the effect of depletion of AP4E1, STX10 or CHMP4C on the overall appearance and activity of the endocytic pathway, uninfected cells depleted of each of these factors were tested for their ability to take up transferrin via the transferrin receptor. Fluorescent transferrin applied to cells is taken up in to recycling endosomes via the transferrin receptor at the cell surface, whereupon it rapidly localises to the endocytic recycling compartment situated around the MTOC (Fig. S2A), demonstrating both the efficiency and kinetics of endosomal uptake. Importantly, this compartment is close to but does not overlap with the TGN (Fig. S2B), indicating that these membranes are separate from this late secretory compartment. In line with its apparent role in TGN-to-plasma membrane transport, depletion of AP4E1 had no effect on transferrin uptake compared to the control siRNA transfected cells (Fig. 4B, Tf uptake), a result that correlated with the comparative CD71 staining of AP4E1 depleted and control transfected cells (Fig. 4B, uninfected, CD71). By contrast, depletion of STX10 had a profound effect on Tf uptake compared to control cells (Fig. 4B, Tf uptake, STX10). This reduction in uptake correlated with a greatly reduced level of transferrin receptor at the cell surface (Fig. 4B, Uninfected, CD71) in line with a previous study of STX10 depleted cells [34]. In the case of CHMP4C depletion, transferrin uptake was altered compared to control cells, with transferrin positive structures distributed throughout the cytoplasm and at the cell surface, (Fig. 4B, Tf uptake, CHMP4C). This alteration reflected a partial reduction in transferrin receptor at the cell surface (Fig. 4B, uninfected, CD71, CHMP4C), indicative of an altered behaviour of the endocytic recycling pathway in the absence of CHMP4C. The relative levels of cell surface transferrin receptor in uninfected cells was recapitulated in HSV1-infected cells, with the reduction in cell surface CD71 in the absence of CHMP4C being more pronounced in infected cells compared to uninfected cells (Fig. 4B, cell surface CD71, infected). To determine if depletion of CHMP4C had an effect on other stages of the endocytic pathway, cells that had been transfected with siRNAs for CHMP4C were also stained for the lysosomal marker LAMP2, the multivesicular/late endosomal marker CD63, or the TGN-to-endosome marker mannose 6 phosphate receptor (M6PR). In all cases, there was no change in the appearance of these compartments when CHMP4C was depleted (Fig. S3), indicating that CHMP4C depletion specifically alters the early and not the late endocytic pathway.

Collectively, these data indicate that CHMP4C depletion is alone amongst the five chosen factors in specifically blocking a very late stage of HSV1 infection, downstream of virus entry, genome replication, protein synthesis and glycoprotein delivery to the plasma membrane. However, CHMP4C is one of three members of the CHMP4 family of proteins [35], and it has been suggested that all three members of this family may have a role in HSV1 morphogenesis [36]. To confirm that CHMP4C and not 4A or 4B are involved, as was implied from the initial screen, HeLa cells were transfected with siRNA for either CHMP4A, CHMP4B, or CHMP4C, either alone or in combination, and after 48 h, infected with virus. After 24 h infection, extracellular virus was harvested and quantified by plaque assay (Fig. S4A). Depletion of CHMP4A or CHMP4B, or CHMP4A and CHMP4B in combination, did not reduce the amount of extracellular virus produced in comparison to cells transfected with the control siRNA. Further, the reduction in amount of virus produced by CHMP4C depleted cells was not enhanced by depleting CHMP4A or CHMP4B in combination. RT-qPCR on HeLa cells that were depleted of either CHMP4A, CHMP4B, or CHMP4C, alone or in combination, was used to determine if transfection with multiple siRNAs affected the efficiency of siRNA knockdown (Fig. S4B). In all cases, transfection of the indicated siRNA resulted in a more than 90% decrease in transcript levels relative to the siRNA control. Altogether, this suggests that only depletion of CHMP4C, and not CHMP4A or CHMP4B, has an effect on HSV1 infection in our experimental system.

### CHMP4C localises to recycling endosomes

CHMP4C is a component of the ESCRT-III fission machinery that would be expected to be involved in membrane scission at various locations in the cell. It has been well-characterised for its role in the abscission checkpoint during cytokinesis, where it localises to the midbody ring to delay cytokinesis and protect against DNA damage accumulation [28, 37]. Although less detail is available on its localisation in the interphase cell, by extrapolation with other ESCRTIII proteins and their role in multivesicular body (MVB) biogenesis, where they play a role in the involution and scission of membranes budding into the lumen of these structures (reviewed in [38]), it would be expected that at least a proportion would localise to the late endocytic pathway. Given the unexpected effect of CHMP4C depletion on recycling endosomes as noted above, its cellular localisation in relation to recycling and late endocytic compartments was assessed next. Endogenous CHMP4C was not detectable with antibodies available to us, so we therefore examined cells expressing V5-tagged CHMP4C (with expression confirmed by Western blotting, Fig. S1A), focusing on cells expressing lower levels of the protein.

Unexpectedly, no colocalization with the MVB marker CD63 (Fig. 5A) was found, but substantial overlap was observed with the transferrin receptor marker for recycling endosomes in cells at different stages of the cell cycle (CD71, Fig. 5B, C & D). First, as described by many others, overexpressed CHMP4C localised to the midbody of cells in the process of cytokinesis (Fig. 5B, arrowed) confirming that it localises as expected when expressed exogenously [28]. In the same cells, a population of CHMP4C localised around the centrosomes and in tubules emanating from the centrosome. Likewise, recycling endosomes were also localised around the centrosomes together with the intercellular bridge, where they are known to deliver membrane to the cleavage furrow [39]. In interphase cells, there was also a high degree of overlap between the CHMP4C signal and the transferrin receptor signal (Fig. 5C) which was particularly obvious in cells that were judged to be entering interphase shortly after cytokinesis (Fig. 5D).

**Figure 5.**
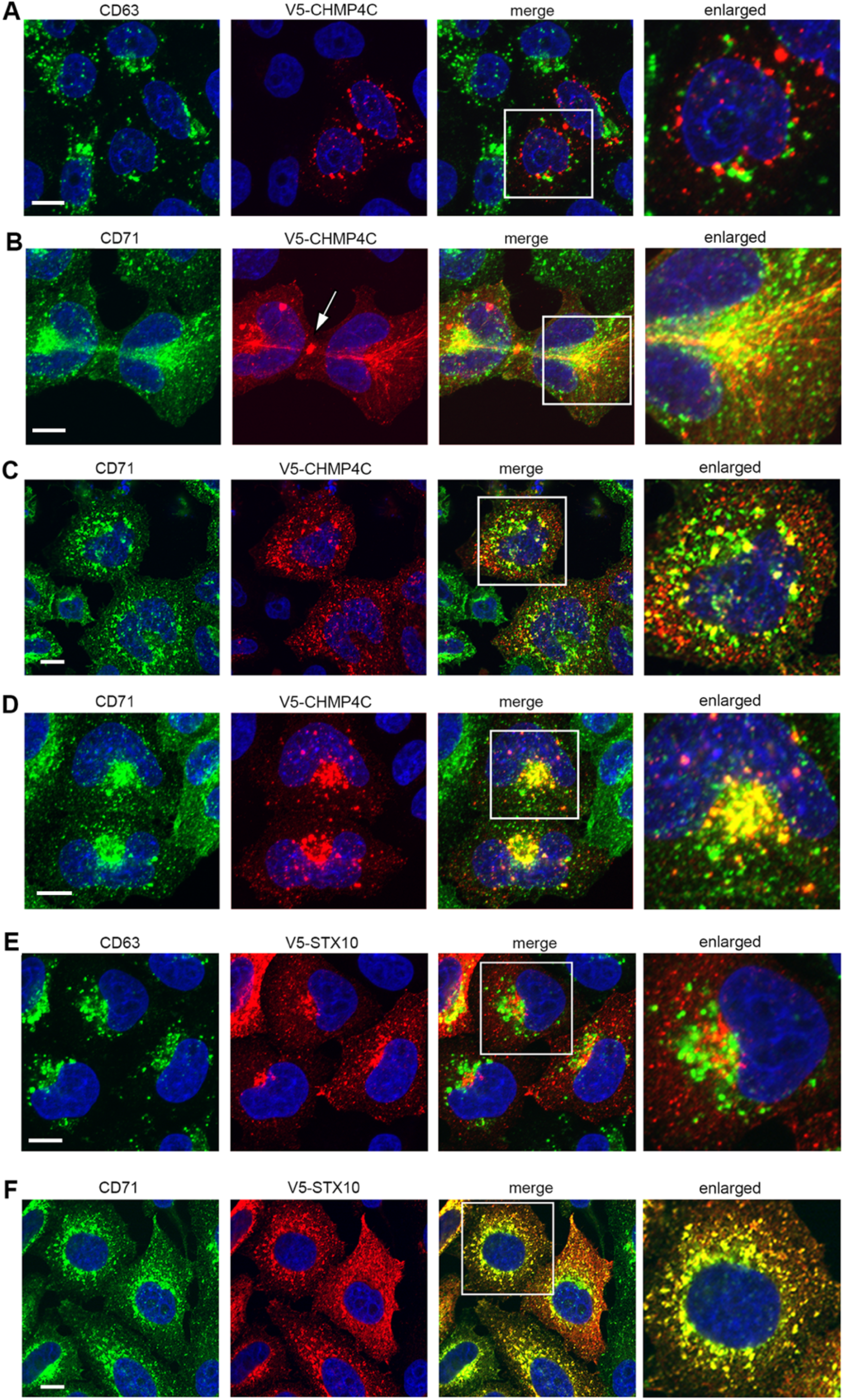
Exogenously expressed CHMP4C and STX10 colocalise with the transferrin receptor. HeLa cells were transfected with plasmid expressing V5-CHMP4C (**A, B, C & D**) or V5-STX10 (**E & F**) and after 24h were fixed and stained for V5 (red) and CD63 (green in **A & E**) or CD71 (green in **B, C, D & F**). Nuclei were stained with DAPI (blue). Scale bar = 10 μm.

Given the effect of STX10 depletion on CD71 localisation to the plasma membrane (Fig. 2B) and its effect on virus replication, we also tested the localisation of exogenously expressed V5-STX10 (confirmed by Western blotting, Fig. S1A). As for CHMP4C, there was no overlap between V5-STX10 and CD63 (Fig. 5E) but there was almost total colocalisation with the transferrin receptor (Fig. 5F), suggesting that this SNARE protein localises to recycling endosomes. Although it has previously been published that STX10 localises to the TGN [26], our results showing co-localisation with the transferrin receptor help to correlate the localisation of this SNARE with its previously identified role, both here and elsewhere, in recycling of the transferrin receptor to the plasma membrane [34].

### Depletion of CHMP4C leads to extensive accumulation and tubulation of endocytic membranes in HSV1-infected cells

As demonstrated above, CHMP4C localises with recycling endosomes and its depletion demonstrably alters the behaviour of the recycling endocytic pathway. Analysis of the transferrin receptor in infected cells revealed that in the absence of CHMP4C, the transferrin receptor exhibited an altered localisation comprising accumulation at the MTOC concomitant with a reduction at the plasma membrane (Fig. 6A), in line with the relative cell surface staining observed above in infected cells (Fig. 4). On closer examination a proportion of cells were found to exhibit long, thin, CD71-positive tubular extensions emanating from the region of the MTOC towards the cell periphery (Fig. 6B, Z projections). These tubules were reminiscent of published studies on brefeldin A (BFA)-treated cells, where it has been shown that BFA, which inhibits Arf GTPases, inhibits coat assembly and subsequent fission of endocytic membranes [40]. Therefore, the staining pattern of CD71 seen in the infected CHMP4C depleted cells was compared to that in BFA-treated cells, fixed at various times after addition (Fig. 6C). Under these conditions the transferrin receptor was found in similarly thin, tubular extensions emanating from the MTOC, as described in previous studies [41].

**Figure 6.**
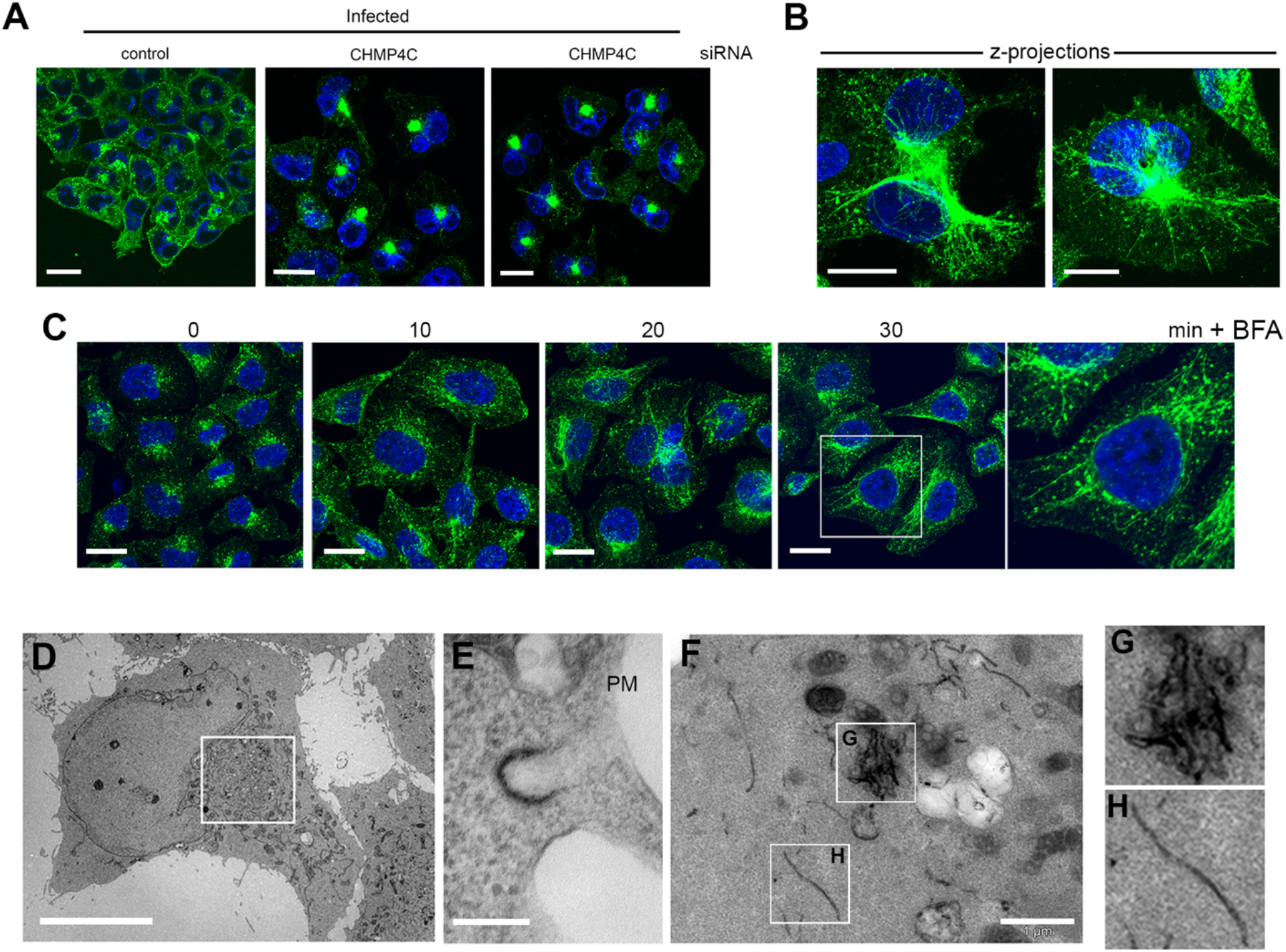
Depletion of CHMP4C leads to tubulation and accumulation of recycling endocytic membranes in HSV1 infection. (**A**) HeLa cells transfected with CTRL or CHMP4C siRNAs were infected with Sc16 for 12 h, fixed and stained for transferrin receptor (CD71). Scale bar = 20 μm. (**B**) As for (**A**), but individual CHMP4C depleted cells are presented as Z-projections of the entire cell volume. Scale bar = 20 μm. (**C**) HeLa cells were treated with BFA (1 μg/ml), fixed at the indicated times thereafter and stained for the transferrin receptor (CD71). Scale bar = 20 μm. (**D**) TEM images of CHMP4C depleted, HSV1 infected cells, showing membrane accumulation in the centre of cells. Scale bar = 10 μm. (**E to H**) As for D but cells were incubated with HRP-transferrin for 30 min prior to fixation and processing for TEM. (**E**) Section (120-nm deep) showing clathrin-coated pit labelled with HRP-transferrin. Scale bar = 200 nm. (**F to H**) Sections (300 nm-deep) showing HRP-transferrin positive tubules in cytoplasm. Scale bar = 1 μm.

Ultrastructural analysis by transmission electron microscopy (TEM) revealed massive membrane accumulations in the cytoplasm next to the nucleus of many infected cells depleted of CHMP4C (Fig. 6D). To further correlate with the confocal microscopy of recycling endosomes, TEM of CHMP4C-depleted, HSV1-infected cells that had been incubated with HRP-transferrin to specifically label transferrin receptor-positive, recycling membranes was carried out. In these cells HRP-transferrin positive clathrin-coated pits were identified (Fig. 6E) indicating the binding of transferrin to its receptor. Sectioning to a depth of 300 nm revealed the presence of extensive, HRP-transferrin positive tubules either clustered in the middle of the cell (Fig. 6F &G) or individually running through the cytoplasm (Fig. 6F & H), confirming that these tubules formed by the depletion of CHMP4C were indeed a component of the transferrin-receptor positive recycling endocytic network. This perturbation of the recycling endocytic network was more pronounced in infected cells than that seen in uninfected cells in the absence of CHMP4C.

### Depletion of CHMP4C inhibits envelopment of HSV1

To investigate how this effect of CHMP4C depletion on the endocytic network affects virus morphogenesis, we used transmission electron microscopy to examine CHMP4C-depleted, HSV1-infected HeLa cells at 16 hpi compared to control siRNA-transfected cells. In control and CHMP4C-depleted infected cells, newly assembled capsids were found inside the nucleus and budding through the nuclear membrane, confirming that capsid assembly occurs as normal in the absence of CHMP4C (Fig. 7, compare A with B & C). By contrast, while many enveloped virions were found released from the control transfected cells (Fig. 7D), in CHMP4C depleted, infected cells, only a small number of enveloped virions were found outside the cell (Fig. 7E), in accordance with the results of our released virus quantification (Fig. 1A; Fig. 4A). Further, unlike the fully enveloped virions ordinarily found wrapped in double membranes in the cytoplasm of HSV1-infected cells [9], in CHMP4C-depleted infected cells, these fully wrapped capsids were much rarer and difficult to detect, despite the presence of numerous capsids within the cytoplasm (Fig. 7F & G, arrowhead). Moreover, many capsids were found associated with incomplete double membranes, suggesting a failure to seal the membrane around the capsid (Fig. 7G, arrowed). Taken together, these results indicate that the depletion of the fission factor CHMP4C alters the integrity of the recycling endocytic network leading to a global accumulation of membranes at the recycling compartment and abrogation of the final wrapping events in HSV1 morphogenesis. In short, this study identifies a new role for CHMP4C in the biogenesis of recycling endosomes.

**Figure 7.**
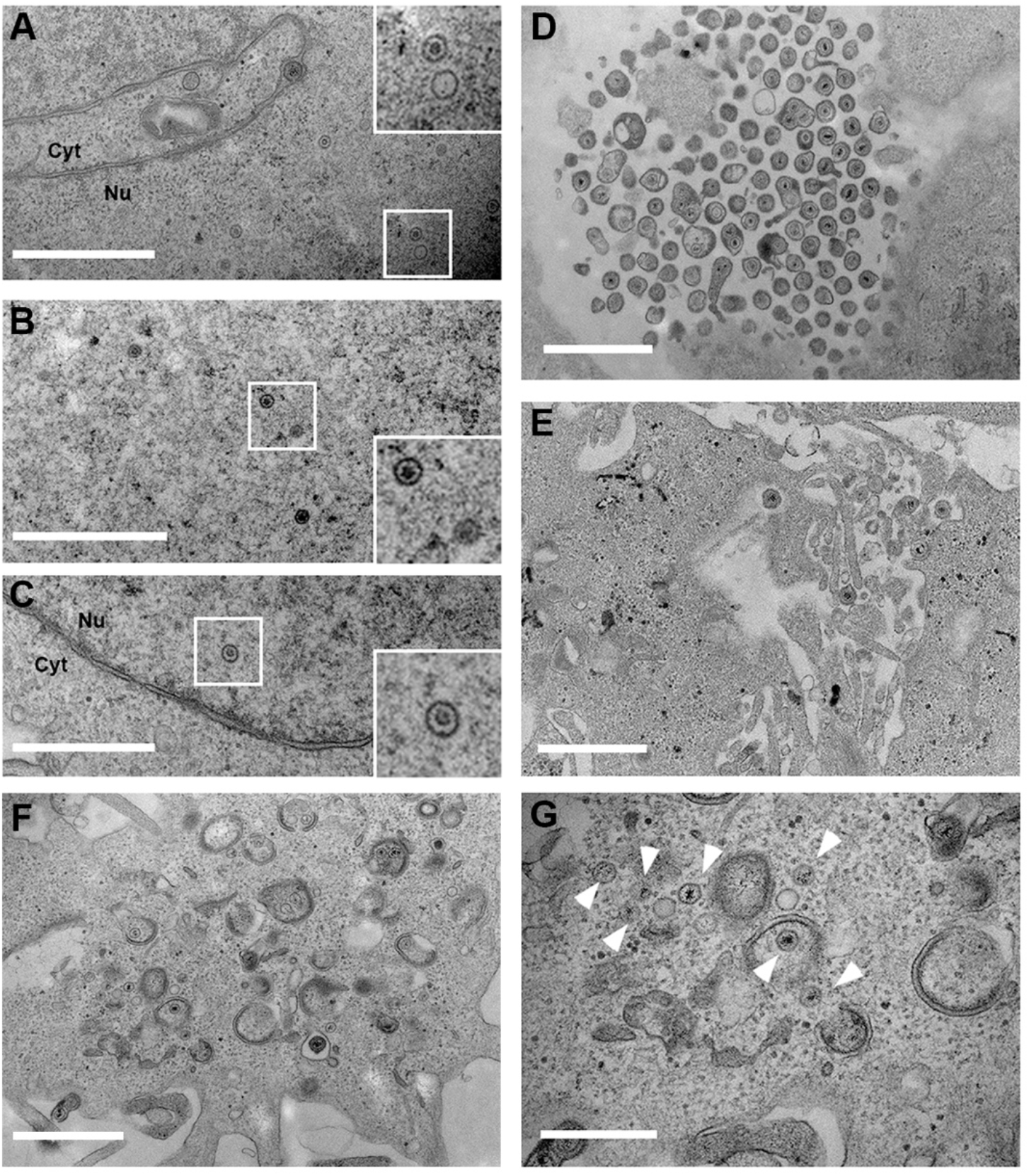
HeLa cells transfected with CTRL (**A & D**) or CHMP4C (**B, C, E, F & G**) siRNAs were infected after 48 h with Sc16 at an MOI 2, and 12 h later were fixed and processed for imaging by TEM. Scale bar = 1 μm (**A** to **F**) or 500 nm (**G**). Arrowheads – capsids in cytoplasm.

## Discussion

The basic replication strategies of viruses, and their exploitation of the host cells they infect, must be understood before novel therapeutics can be developed. Many highly pathogenic human viruses are surrounded by lipid envelopes that are rich in virus-encoded proteins, and are derived from intracellular membranes. These pathogens are therefore highly dependent on cellular trafficking pathways for the production of infectious virus. Indeed, many potentially share common routes of envelopment, raising the possibility of targeting envelopment as a broad-spectrum intervention strategy. This study focused on the alphaherpesvirus HSV1, a large enveloped DNA virus, which has a complex morphogenesis pathway involving the nucleus, the cytoplasm and many aspects of cellular membrane trafficking pathways [42]. Given our dual goals to understand virus manipulation of cellular trafficking pathways and to identify points of possible intervention, we chose to target a rationally selected set of cellular trafficking factors rather than conducting a genome-wide screen. As proof of principle, all human Rab GTPases were screened previously for involvement in HSV1 morphogenesis, with four (Rabs 1, 6, 5 and 11) identified as involved in the trafficking of virus glycoproteins and subsequent envelopment of the virus in endocytic tubules [9, 15]. In this current study a range of 82 cellular trafficking factors covering different functions in the secretory and endocytic pathways were targeted, and eleven were identified whose depletion reduced infectious virus production by > 20-fold, with an additional 14 reducing virus production by > 10-fold, indicating a potential role for these proteins in HSV1 infection. This outcome compares favourably with a recent study on the betaherpesvirus hCMV, where a screen of 156 host factors involved in membrane organisation found that depletion of 15 factors reduced virus yield between 5- and 12-fold [43]. Moreover, a number of genome-wide RNAi screens of different enveloped virus infections have all identified around 300 targets out of ~ 20,000 genes tested [44–47], indicating that our targeted screen was efficient in discovering important factors.

Of the five trafficking factors that were investigated in detail, two were found to inhibit the virus life-cycle at early stages of infection prior to morphogenesis. First, depletion of the COPG1 coatamer subunit γ1, which is one of seven subunits that form the stable coat complex COPI [23], was found to have the most profound effect on virus production in both library screens. COPI coats form on vesicles and tubules and are responsible for retrieval of proteins from the Golgi to the ER [48]. COP1 also localises to endosomes, but its role there is unclear [49]. Not surprisingly, COPI appears to be exploited by many viruses throughout their lifecycles [50], and has recently been shown to be required for hCMV infection using a similar siRNA screen as described here [43]. Nonetheless, although a potential role for COPG1 downstream in virus infection/trafficking of virus glycoproteins through the secretory pathway cannot be ruled out, data presented here show that the predominant effect of COPG1 depletion is the inhibition of virus entry, potentially by reducing the level and presentation of the HSV1 receptor, nectin1, at the plasma membrane. It should also be noted that COPG1 depletion affected the viability of cells which in turn could reduce their ability to support early stages of virus infection (Fig. 1D). Second, depletion of the v-SNARE protein VAMP4 had a profound effect on HSV1 genome replication, upstream of virus morphogenesis, and as a consequence VAMP4-depleted cells supported a reduced level of late protein synthesis. Given that VAMP4 is enriched in the TGN and cycles from the cell surface to the TGN [27], this may suggest that rather than directly affecting glycoprotein trafficking, its depletion results in a global perturbation of cell integrity similar to that seen for COPG1 depletion, as evidenced by the reduced viability of VAMP4-depleted cells (Fig. 1D). This global effect may therefore specifically affect the process of genome trafficking or genome replication in the nucleus. In short, while both these factors have important roles in intracellular trafficking, neither can be assigned a direct role in virus morphogenesis.

Three of the factors that were investigated were shown to perturb HSV1 morphogenesis specifically. AP4E1 is a late secretory pathway protein, and is a component of the adaptor protein complex AP4, a poorly characterised complex involved in non-clathrin coat vesicle formation and cargo selection on vesicles departing the TGN [51]. AP4 is present at a relatively low abundance but is ubiquitously expressed, and has been proposed to be involved in trafficking of specialized cargoes such as the transport of ATG9 to autophagosomes [52, 53]. Although speculative at this stage, it is possible that HSV1 utilises AP4 to transport one or more of its glycoproteins out of the TGN, which may explain the reduced localisation of gD and gE at the cell surface in AP4E1 depleted infected cells (Fig. 4A), and further work will be required to determine if any glycoproteins contain functional AP4-binding motifs. The final two factors of the five tested, CHMP4C and STX10, were both found to localise to and be required for the integrity of recycling endosomes. This was unexpected for STX10 which has been shown previously to localise to the TGN [26, 34]. Nonetheless, it is also required for recycling the transferrin receptor to the plasma membrane [34], a result that was confirmed here, placing its role within the recycling endocytic network and suggesting that its co-localisation with the transferrin receptor in recycling endosomes may be functionally relevant. CHMP4C on the other hand is a component of the ESCRT-III fission machinery involved in membrane remodelling during various processes within the cell, including late endosomal sorting, plasma and nuclear membrane repair, and abscission during cytokinesis [38]. However, it has not been previously located on recycling endosomes, or shown to be required for their biogenesis. Amongst the other factors identified as reducing virus yield and not examined further (Table 1) the coat protein clathrin light chain A, while not being required for the formation of clathrin coats at the plasma membrane or the TGN, is required for coat formation in dynamic tubular endosomes involved in recycling to the plasma membrane [54]. Additionally, dynamin, which was also identified as a hit in the second siRNA screen is involved in the recycling of the transferrin receptor to the plasma membrane by the aforementioned endosome-derived clathrin-coated vesicles [55]. Collectively, these results identify multiple factors involved in the biogenesis of recycling endosomes that are important for HSV1 infection, adding weight to the importance of the recycling endocytic network in the morphogenesis of HSV1.

CHMP4 is the most abundant component of the ESCRT-III membrane-remodelling machinery, and has a role in the envelopment of human immunodeficiency virus (HIV) [56], along with other viruses such as dengue virus [57]. While there are three paralogues of CHMP4, the efficient depletion of CHMP4A or CHMP4B in isolation or in combination had no effect on the level of HSV1 production, indicating that the perturbation of HSV1 morphogenesis was highly specific to CHMP4C depletion (Fig. S3). This is in contrast to the situation with HIV budding where CHMP4B but not A or C is required [56]. Nonetheless, our results differ from a previous study on HSV1 that did not deplete but used overexpressed dominant-negative CHMP4 proteins to show that all three dominant-negative CHMP4 paralogues interfered with HSV1 production, with a greater effect of CHMP4B overexpression compared to the others [36]. A major role of CHMP4C in the cell has been defined as a checkpoint protein to prevent chromatin mis-segregation during abscission in cytokinesis [28]. During cytokinesis, CHMP4C localises to the midbody whereas CHMP4B and A localise on either side of the midbody in preparation for membrane scission. Indeed, depletion of CHMP4C accelerates cytokinesis due to checkpoint inactivation, whereas depletion of CHMP4B results in a failure of abscission [28]. Hence, our discovery that CHMP4C but not A or B is required in HSV1 morphogenesis identifies another role for CHMP4C that separates it from the other CHMP4 paralogues.

Data presented here suggest that in at least some situations, CHMP4C is involved in the fission of tubular recycling endosomes. While less pronounced in uninfected cells, recycling endosomes were aberrantly localised to the MTOC in HSV1-infected cells, with thin tubules emanating towards the periphery of the cell, indicating a perturbation to the normal trafficking and biogenesis of this compartment. Given that this network is known to be important in HSV1 envelopment [9], the profound effect of CHMP4C depletion together with HSV1 infection on the appearance of these membranes may indicate that HSV1 specifically activates and utilises a population that require CHMP4C for their correct scission. Extensive tubulation of recycling endosomes has been detected in other related scenarios, such as BFA-mediated inhibition of Arf GTPases, required for coat formation and budding from membranes [40]; knockdown of endosome-localised Arf GTPases themselves [58]; and knockdown of the BIG2 guanidine exchange factor (GEF) for such Arfs [59]. Although it is not yet known if this tubulation represents abrogated fission of these membranes on the way to the ERC from sorting endosomes, or on the way out of the ERC on the way to the plasma membrane [60], it is clear that the tubulation/accumulation phenotype represents a defect in the biogenesis of recycling endosomes. Moreover, the capsid wrapping profiles that were detected in CHMP4C-depleted infected cells, whereby the capsid was frequently observed in association with a cup-like double membrane that had not yet sealed to form a double enveloped virion (Fig. 7H) may reflect a requirement for CHMP4C in the closure of the neck of this structure, a process that would share topology with canonical ESCRT-III dependent processes [38]. Indeed, one of the identified roles of ESCRT-III is the sealing of the phagophore in autophagy [61], a process that is similar at the physical level to our working model of HSV1 envelopment [9]. Of note, CHMP2A, which was shown recently to regulate phagophore closure [62], was also picked up here in our second library screen as a possible factor involved in HSV1 infection (Table 1).

It is becoming increasingly accepted that host cell processes required for virus infection could be appropriate targets for antiviral intervention [63, 64], and virus morphogenesis and envelopment may offer an opportunity for such intervention. Moreover, the study of virus morphogenesis provides the opportunity to identify functions of as yet uncharacterised cellular trafficking proteins. Our discovery of a new role for CHMP4C in the biogenesis of HSV1-wrapping membranes paves the way for future work in both areas.

## Materials and Methods

### Cells and viruses

Vero and HeLa cells were cultured in Dulbecco’s modified Eagle’s medium (DMEM) supplemented with 50 U/mL penicillin/streptomycin and 10% foetal bovine serum (FBS). All viruses used were routinely propagated in Vero cells in DMEM supplemented with 2% newborn calf serum (NCS) and 50 U/mL penicillin/streptomycin. All plaque assays were carried out in Vero cells in DMEM supplemented with 2% NCS, 50 U/mL penicillin/streptomycin and 1% human serum (BioIVT). HSV1 strain Sc16 was used routinely. Sc16 110lacZ [65] has been previously described and was provided by Stacey Efstathiou (University of Cambridge).

### siRNAs and transfections

Silencer select siRNA duplexes (Ambion, ThermoFisher Scientific) were forward or reverse transfected with Lipofectamine 2000 (Invitrogen) in HeLa cells to a final concentration of 20 nM according to the manufacturer’s instructions. The Silencer Select negative control siRNA number 1 was used as a negative control (Ambion, ThermoFisher Scientific).

### Plasmids

CHMP4C and STX10 open reading frames were amplified by PCR from HeLa cDNA using primers shown in Table S5 and inserted into pCR-BluntII-TOPO cloning plasmid. CHMP4C and STX10 open reading frames were transferred as BamH1-EcoR1 and BamH1-BamH1 fragments respectively into an in-house expression vector driven by CMV-IE promoter [66], which places the V5 tag at the N-terminus of both proteins. Nectin1A was first amplified from HeLa cDNA using primer set 1 shown in Table S5 and inserted into pCR-BluntII-TOPO cloning plasmid. The nectin1A open reading frame was subsequently amplified by PCR using primer set 2 and inserted into pEGFPN1 as a BamH1-Not1 fragment, incorporating the V5 epitope tag at the C-terminus of nectin1A.

### Antibodies and reagents

The following primary antibodies were used for Western blots and immunofluorescence, and were kindly provided by: mouse anti-gD (LP14) and mouse anti-gB (R69), Tony Minson (University of Cambridge); mouse anti-VP16 (LP1), Colin Crump (University of Cambridge); mouse anti-gE, David Johnson (Oregon Health and Science University); mouse anti-nectin1 (CK6), Claude Krummenacher (Rowan University, New Jersey). Our rabbit VP22-specific antibody (AGV031) has been described elsewhere [67]. Commercially available antibodies using in this study included: mouse anti-V5, mouse anti-gC, mouse anti-β actin, mouse anti-CD63 and rabbit anti-giantin (all AbCam); mouse anti-CD71 (SantaCruz); mouse anti-γ tubulin and mouse anti-a tubulin (Sigma). Goat anti-mouse IRDye 680RD and goat anti-rabbit IRDye 800CW (LI-COR Biosciences) secondary antibodies were used as appropriate for Western blots, while AlexaFluor conjugated secondary antibodies (Invitrogen) were used for immunofluorescence. Endocytic structures were labeled by incubating cells with texas red conjugated transferrin (Invitrogen), Alexa 488 conjugated transferrin (Molecular Probes) at a concentration of 1 μg/ml for 30 min. Brefeldin A (Sigma) was used at a concentration of 1 μg/ml.

### Viability assay

Cell viability was assessed using a CellTitre-Glo luminescent viability assay (Promega) and read on a CLARIOstar microplate reader (BMG Labtech).

### SDS-PAGE and Western blots

Samples were separated by SDS-PAGE (10 – 14% polyacrylamide as appropriate) and transferred to nitrocellulose membranes before blots were imaged using an Odyssey CLx imaging system (LI-COR Biosciences).

### *N*-linked glycosylation analysis

Cell lysates were lysed on ice in glycosylation lysis buffer (50 mM Tris base pH 7.5, 200 mM sodium chloride, 2 mM magnesium chloride and 1% v/v NP-40) with protease inhibitor cocktail (ThermoFisher). Glycoprotein denaturation of the cell lysate was performed with 1x glycodenaturing buffer (New England Biolabs) at 95°C for 10 mins. Denatured cell lysate was then deglycosylated using 500 U of PNGase F (New England Biolabs) in 1x glycobuffer 2 (New England Biolabs), and 10% NP-40 (New England Biolabs) at 37°C for 60 mins.

### Transmission electron microscopy

To prepare samples for electron microscopy, cells were grown to approximately 50% confluency before forward transfection with siRNA duplexes to a final concentration of 20 nM. After 48 h, cells were then infected with HSV1 Sc16 at a multiplicity of infection of 5 for 12 h. Endocytic events were labelled by incubating cells with medium containing transferrin-HRP (Jackson ImmunoResearch) at a concentration of 1 μg/ml for 30 min prior to fixation in 0.5% glutaraldehyde in 200 mM sodium cacodylate buffer for 30 min before being washed in sodium cacodylate buffer. After fixation, samples were washed and stained with a metal-enhanced DAB substrate kit (ThermoFisher Scientific). The samples for electron microscopy were fixed and processed as described previously [32].

### RNA isolation, reverse transcription and qPCR

RNA was isolated from cells using a RNeasy mini kit (Qiagen), and then 400 ng to 1 μg of RNA was DNase I-treated according to the manufacturer’s protocol (ThermoFisher Scientific). Superscript III (Invitrogen) was used with random primer mix to make cDNA according to the manufacturer’s instructions in a Veriti 96-well Thermal Cycler (Applied Biosystems). All quantitative polymerase chain reactions (qPCRs) were assembled using the MESA BLUE qPCR kit for SYBR assay (Eurogentec) according to the manufacturer’s instructions, with the primer sets listed in Table 1. The qPCRs were carried out on a LightCycler96 system (Roche).

### Quantification of viral DNA

HeLa cells were reverse transfected with 20 nM siRNA duplexes and incubated for 48 h, before infection with HSV1 Sc16 at an MOI of 2, including a control of untransfected cells infected in the presence of 100 ng/ml cytosine arabinoside (AraC). After 1 h, cells were subjected to a gentle acid wash to inactivate any virus that had not entered the cells. After a total of 2 or 24 h infection, DNA was harvested using the DNeasy blood and tissue kit (Qiagen). The qPCR assays were carried out in a LightCycler96 system (Roche), using MESA BLUE qPCR kit for SYBR assay (Eurogentec) according to the manufacturer’s instructions with primers for 18S (see Table 1) and HSV1 *UL48* gene.

### Immunofluorescence

Cells for immunofluorescence were grown on coverslips and fixed with 4% paraformaldehyde in PBS for 20 min at room temperature, followed by permeabilisation with 0.5% Triton-X100 for 10 min. Fixed cells were blocked by incubation in PBS with 10% NCS for 20 min, before the addition of primary antibody in PBS with 10% NCS, and a further 30-min incubation. After extensive washing with PBS, the appropriate Alexafluor conjugated secondary antibody was added in PBS with 10% NCS and incubated for a further 30 min. The coverslips were washed extensively in PBS and mounted in Vectashield (Vector Labs) containing DAPI. Images were acquired using a Nikon A1 confocal microscope and processed using ImageJ software [68].

## Acknowledgements

Thanks to Stacey Efstathiou, Tony Minson, Colin Crump, David Johnson and Claude Krummenacher for kindly providing reagents used in this study.

This work was funded by a project grant to GE from the Medical Research Council (MR/M020061/1). JK was funded by a British Skin Foundation studentship (035/s/17). GLS was supported by a Wellcome Trust Principal Research Fellowship (grant 090315).

## Supplementary Material

**Table S1.** Genes for human trafficking factors targeted by siRNA libraries.

**Table S2.** Relative virus yield from siRNA (set 1) library transfected cells in three independent experiments.

**Table S3.** Relative virus yield from siRNA (set 2) library transfected cells in three independent experiments.

**Table S4.** RT-qPCR primers used for measuring siRNA knockdown efficiency.

**Table S5.** PCR primers for amplification and cloning from HeLa cell cDNA. Bold, underlined sequences are restriction sites used for cloning. Lower case is V5 epitope sequence.

**Figure S1.** Transient expression of V5-tagged constructs used in this study. (**A**) HeLa cells were transfected with plasmids expressing V5-CHMP4C, V5-STX10 or nectin1-V5, and analysed 16 h later by SDS-PAGE and Western blotting with antibody to the V5 epitope and a-tubulin. (**B**) As for (**A**), but nectin1-V5 transfected cells were harvested and deglycosylated with PNGaseF prior to analysing by SDS-PAGE and Western blotting. (**C**) HeLa cells grown on coverslips were transfected with nectin1-V5-expressing plasmid. Sixteen hours later, cells were either cell-surface stained with antibody to the extracellular domain of nectin1 prior to fixation, or fixed and permeabilised followed by staining with the same antibody (green). Nuclei were stained with DAPI (blue). Scale bar = 20 μm.

**Figure S2.** Uptake of transferrin into HeLa cells. (**A**) HeLa cells were incubated with texas red conjugated transferrin for 30 min, before fixing and staining for γ tubulin (green) to label the MTOC. (**B**) HeLa cells were incubated with FITC-transferrin (green) for 30 mins, before fixing and staining for TGN46 (red). Scale bar = 5 μM.

**Figure S3.** Depletion of CHMP4C has no effect on the late secretory pathway. HeLa cells were transfected with control or CHMP4C siRNAs and fixed and stained two days later for the lysosomal marker LAMP2, the late endosomal marker CD63 or the mannose 6 phosphate receptor M6PR.

**Figure S4.** Depletion of CHMP4C but not CHMP4A or CHMP4B reduces HSV1 production. (**A & B**) 5 nM of each siRNA was reverse transfected alone or in combination as indicated, with controls transfected with equivalent amounts of negative control siRNA. (**A**) After 48 h, cells were infected with HSV1 Sc16 at an MOI 5. After 24 h infection, extracellular virus was harvested and quantified by plaque assay (mean±SEM, *n* =3). (**B**) After 48 h cells were harvested for RNA isolation, and the % siRNA knockdown (kd) was measured for CHMP4A, CHMP4B or CHMP4C by qRT-PCR (mean±SEM, *n* = 3).

